# Investigation of the Effects of N-Linked Glycans on the Stability of the Spike Protein in SARS-CoV-2 by Molecular Dynamics Simulations

**DOI:** 10.1101/2022.01.07.475397

**Authors:** E.Deniz Tekin

## Abstract

We perform all-atom molecular dynamics simulations to study the effects of the N-linked glycans on the stability of the spike glycoprotein in SARS-CoV-2. After a 100 ns of simulation on the spike proteins without and with the N-linked glycans, we found that the presence of glycans increases the local stability in their vicinity; even though their effect on the full structure is negligible.

**Graphical Abstract:** 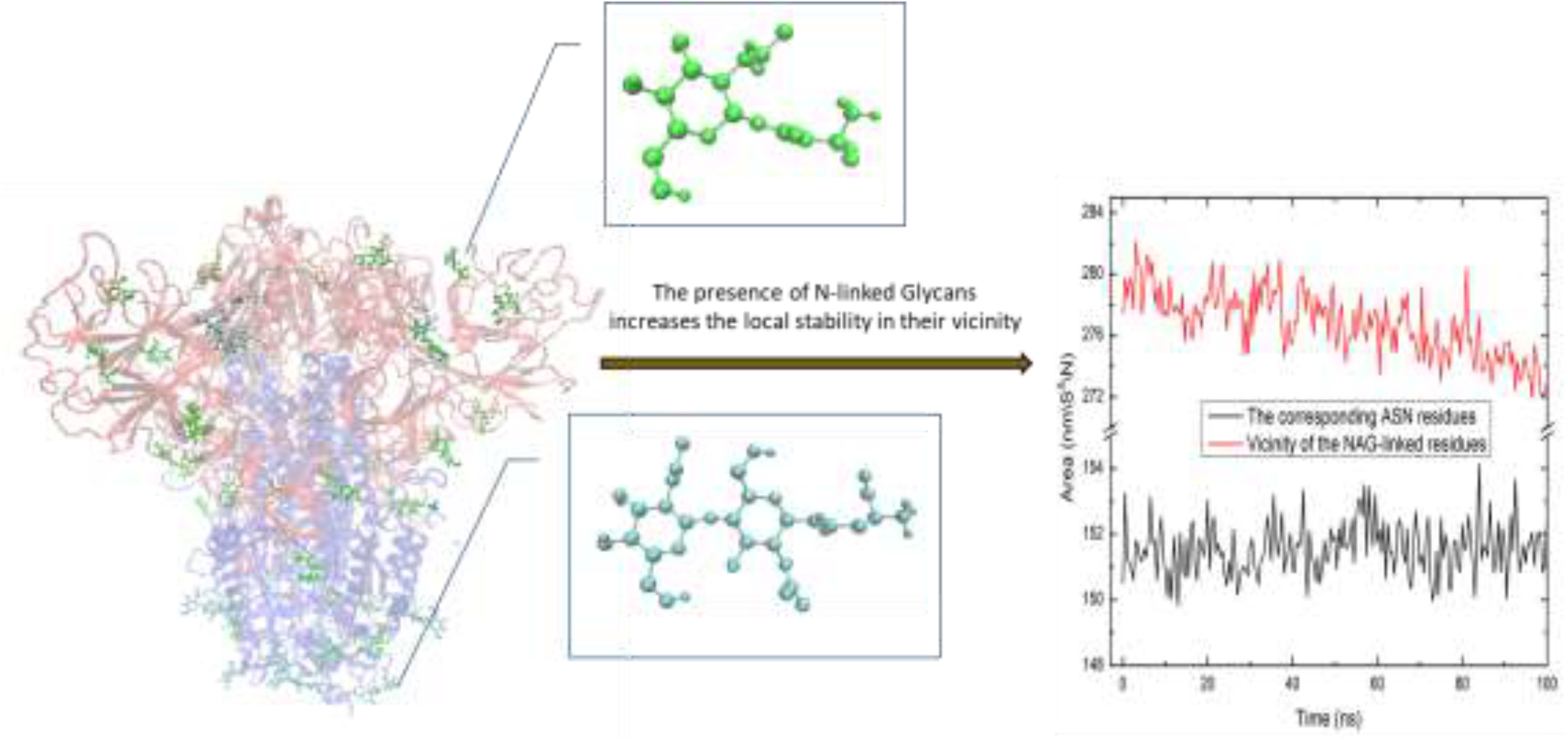

## Introduction

Main structural components of SARSCoV-2 are made up of several different types of proteins: the transmembrane glycoprotein, the spike (S) glycoprotein and the envelope protein; with each protein having definite functions in binding to a receptor and sneaking into the host cell. The largest of these proteins, the S protein, assumed to be the most important one in entering the host cell, is the target of antibodies. For this reason, the three-dimensional shape and dynamics of this protein is extremely important to understand [1]. The SARSCoV-2 S glycoprotein exists as a homotrimer with each monomer composed of two sub-units: the S1 subunit that binds to the host cell receptor, and the S2 subunit that takes part in the fusion of the viral and the cellular membranes. In addition, a furin cleavage site at the boundary between the S1/S2 subunits exists [2]. The boundary, cleaved during the biosynthesis, is a novel feature of this virus which does not exist in the other known coronaviruses such as the SARS-CoV and the SARSr-CoV. These two domains of the S glycoprotein are shown in Figure 1. Walls *et al*. [**1**] determined the cryo-EM structures of the SARSCoV-2 S glycoprotein in two distinct conformations and stored the data as 6VXX.pdb (closed SARS-CoV-2 S) and 6VYB.pdb (SARS-CoV-2 S with one S^B^ open) in RCSB PDB.

**Figure 1.**
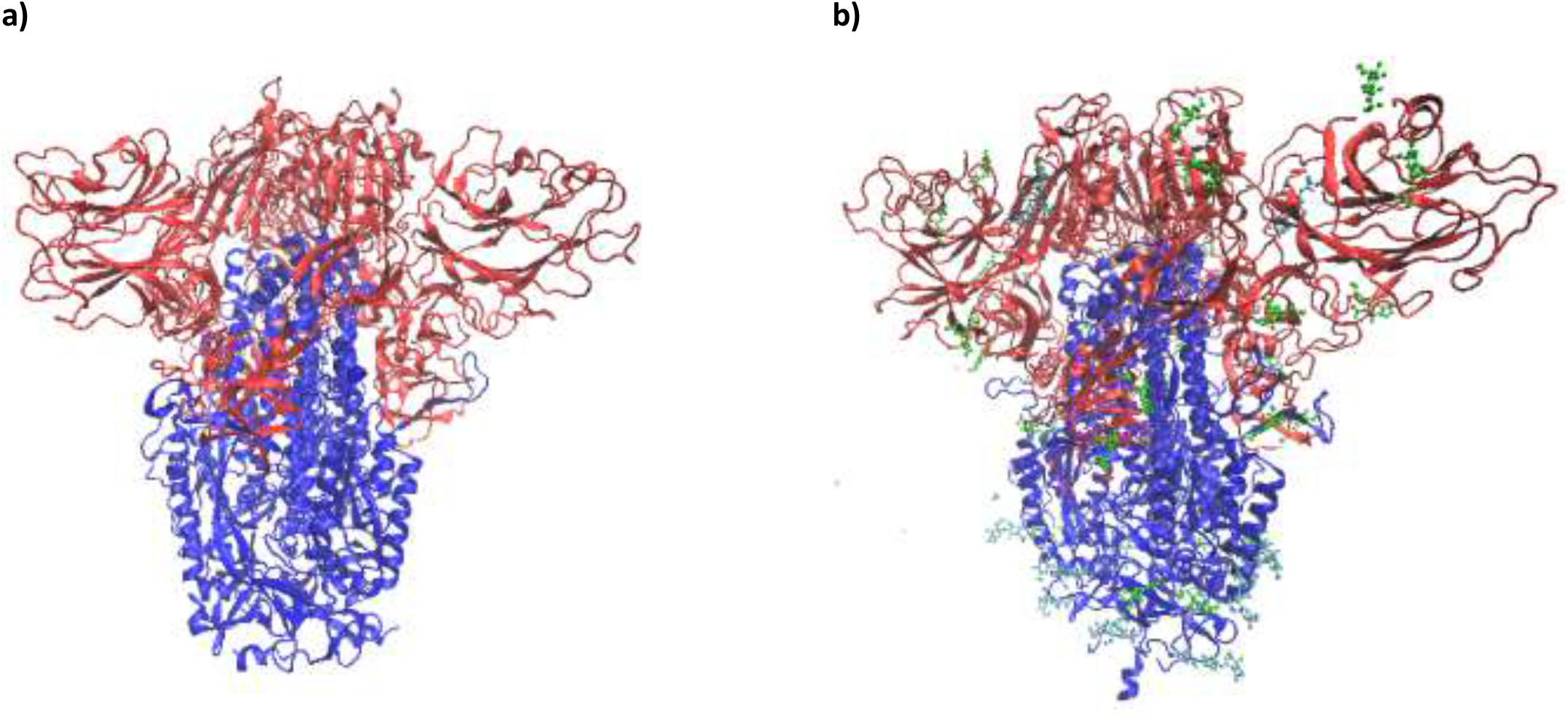
S1 (red) ve S2 (blue) domains of **a)** S-without-NAG **b)** S-with-NAG

Various related coronavirus S glycoproteins are extensively patched with heterogeneous N-linked glycans (NAG) hanging from the trimer surface that are important for proper folding [3] and “for modulating accessibility to host proteases and neutralizing antibodies” [4-6]. Extensive N-linked glycans have also been characterized at the interface of SARS-CoV-2 S (Figure 2); and the existence of glycans is believed to be one of the reasons why the coronavirus has caused a large number of infections and mortality [7-9]. Some comparative studies have been conducted between the coronavirus spikes [10-14] and the coronavirus spikes with other viral proteins [15, 16]. Wang *et al*. summarized the importance of the S protein glycans in viral entry and antibody production in [17]. To date, a large amount of experimental [18-23] and theoretical studies [24-32] have been conducted elucidating the structure of the coronavirus and how it infects. Among them, Amaro *et al*. [33] showed that two N-glycans at positions N165 and N234 have an important structural role in stabilizing the receptor-binding domain (RBD) of S1 “up” conformation using all-atom molecular dynamics simulation (MD). Another MD study conducted by Woods *et al*. [9] revealed the importance of the S protein glycan in shielding the immune recognition. 16 out of 22 SARS-CoV-2 S N-linked glycans (NAGs) per monomer (in total 48) were observed in cryo-EM [1] (Table-1, Figure 2b); however, computer simulation of closed form (6Vxxx.pdb) spike protein with NAG molecules is not easy for the reason that the existing force fields are generally defined for amino acids, nucleic acids and for some special cofactors such as the ATP; but not defined for such mixed molecules. So, I introduced the NAG-linked amino acids to the system as pseudo-amino acids. Details are in the “Material and Methods” section.

**Figure 2.**
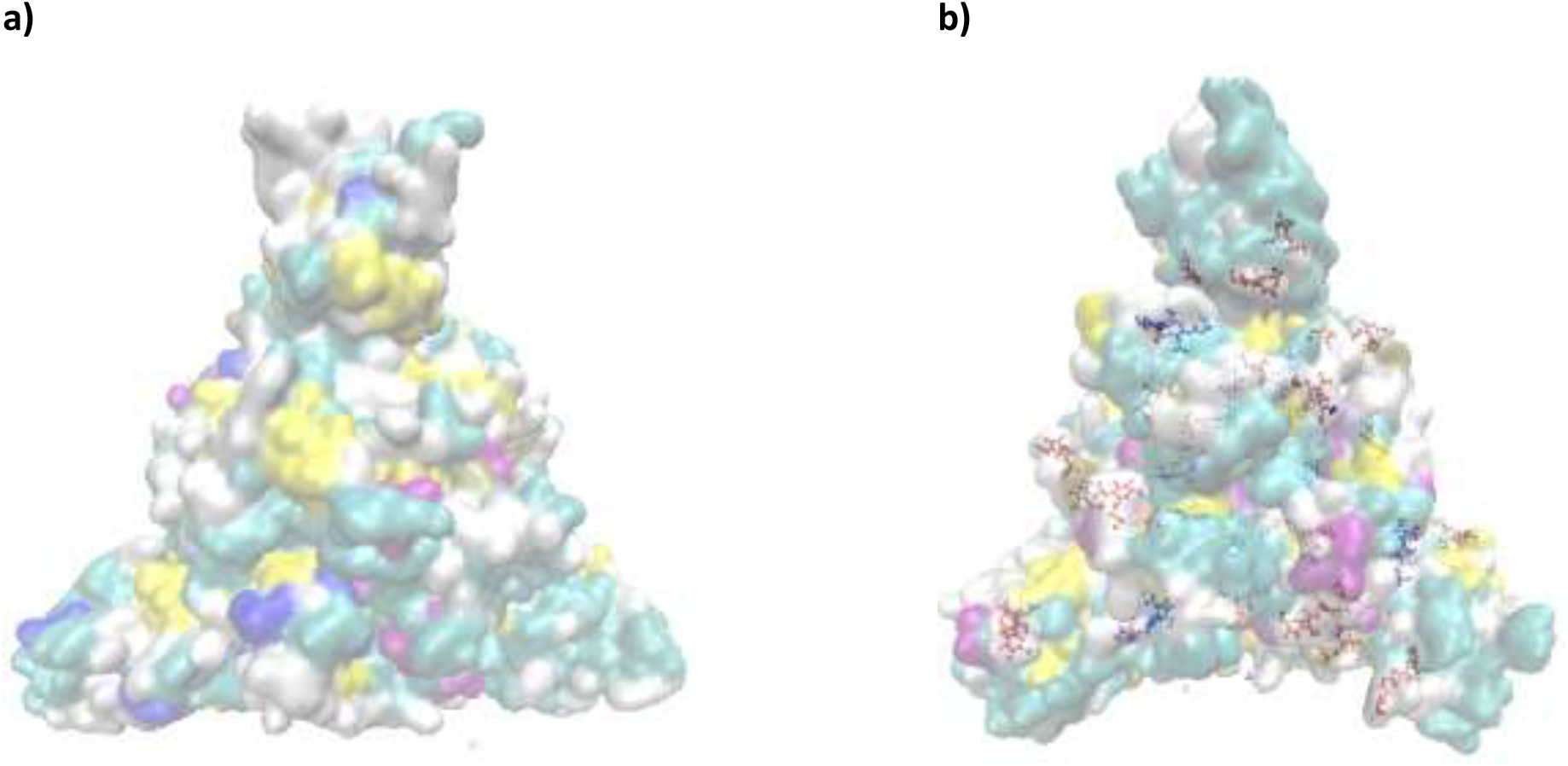
“QuickSurf” representation of SARS-CoV-2 S Protein **a)** without NAG **b)** with NAG at the interface*. *(* N-linked glycans are represented by Blue CPK which we call as ANX and Red CPK which we call as NAN)*

**Table 1:** Pozitions of Glycans in SARS-CoV-2 S.

| N-Linked Glycans<br>(in Ref. [1]) | Corresponding residues / given names<br>(in this simulation) |  |
| --- | --- | --- |
| N<sub>61</sub>VT | N<sub>80</sub>VT | ANX |
| N<sub>122</sub>AT | N<sub>141</sub>AT | ANX |
| N<sub>165</sub>CT | N<sub>184</sub>CT | ANX |
| N<sub>234</sub>IT | N<sub>253</sub>IT | NAN |
| N<sub>282</sub>GT | N<sub>301</sub>GT | ANX |
| N<sub>331</sub>IT | N<sub>350</sub>IT | ANX |
| N<sub>343</sub>AT | N<sub>362</sub>AT | ANX |
| N<sub>603</sub>TS | N<sub>622</sub>TS | ANX |
| N<sub>616</sub>CT | N<sub>635</sub>CT | ANX |
| N<sub>657</sub>NS | N<sub>676</sub>NS | ANX |
| N<sub>709</sub>NS | N<sub>728</sub>NS | ANX |
| N<sub>717</sub>FT | N<sub>736</sub>FT | NAN |
| N<sub>801</sub>FS | N<sub>820</sub>FS | NAN |
| N<sub>1074</sub>FT | N<sub>1093</sub>FT | ANX |
| N<sub>1098</sub>GT | N<sub>1117</sub>GT | NAN |
| N<sub>1134</sub>NT | N<sub>1153</sub>NT | NAN |

In this work, 100 ns MD simulations of closed SARS-CoV-2 S (6VXX.pdb) without NAG (S-without-NAG) and SARS-CoV-2 S with NAG (S-with-NAG) have been performed to better understand the role of the NAG molecule on the stability of the S protein.

## Material and Methods

### Setup of the system

The structure of the SARS-CoV-2 S glycoprotein (closed state) was taken from the Protein Data Bank with the 6vxx code [1]. The missing residues were added using the Swiss Pdb-Viewer program [34] in order to complete the structure before running the molecular dynamics simulations. The acetyl (ACE) and amino (NH2) groups were capped to each -mer of the S protein, in order to keep the protein in a fully folded state. The difficulty one encounters is that the NAG molecule, and so the NAG-linked amino acid, is not a known entity in any force field of GROMACS [35]. In the S protein, NAG is linked to the Asparagine (Asn) in two configurations: Mono-Saccharide NAG bound to ASN which I called as ANX (Figure 3a) and Poly-Saccharide NAG residues bound to ASN which I called as NAN (Figure 3b). To be able to introduce the NAG-linked amino acids, ANX and NAN, into an existing force field, one needs to parametrize them. To this end, geometries of the ANX and NAN were created by ArgusLab [36] and the bonded parameters (bonds, angles, dihedrals and impropers) were added to the aminoacids.rtp file of Gromacs 54a7 force field [37]. The partial charges derived from the electrostatic potential were obtained by single point calculations at the B3LYP-D3/6-31G**++ level of theory using the Jaguar 10.7 program package [38]. Then, ANX and NAN were defined as Protein in the residuetypes.atp file. Thus, we obtained the complete model of the S glycoprotein of SARS-CoV-2 (S-with-NAG). Besides, in order to understand what effects the NAG molecule has on the S protein, we also created a system which is devoid of the NAG molecules (S-without-NAG) for comparison.

**Figure 3.**
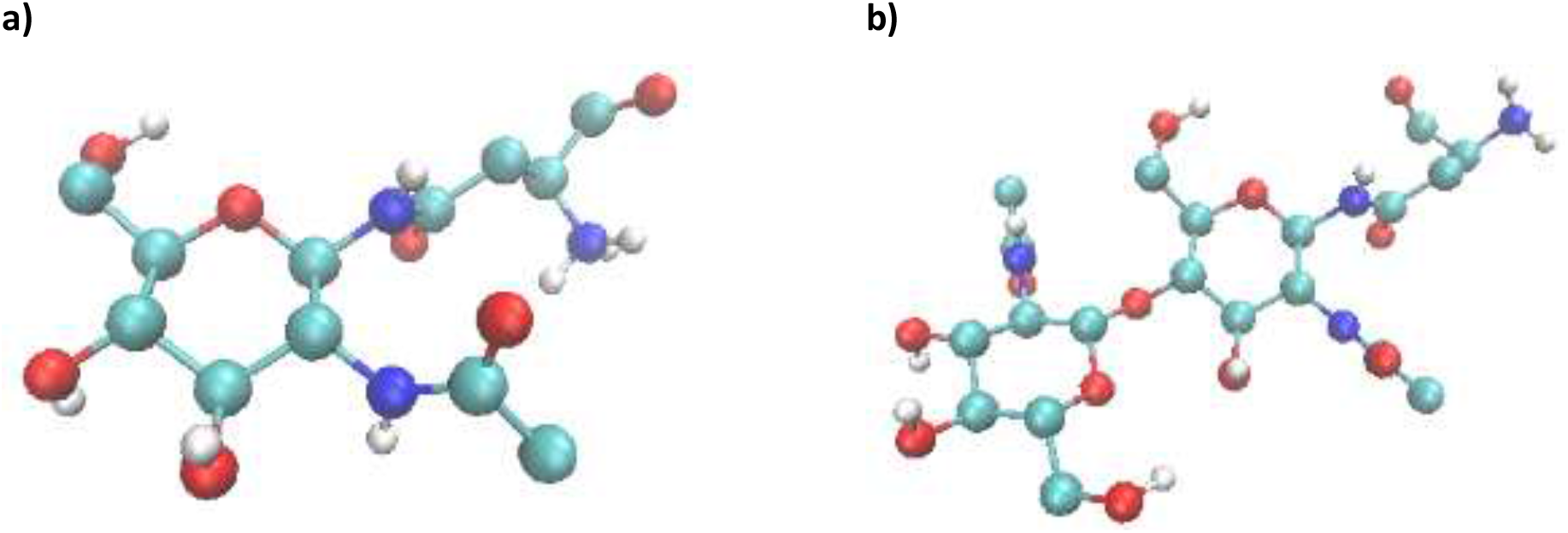
Structures of **a)** ANX and **b)** NAN

### Molecular Dynamics (MD) Simulations

S-without-NAG and S-with-NAG proteins were put in separate rhombic dodecahedron box filled with a SPC [39] type water molecules. Minimum image convention is set with the S-trimers in the center of the boxes and at a distance 1.0 nm to the edges of the boxes. We added 15 Na^+^ ions (5 Na^+^ ions per - mer) to neutralize the charge of the systems. Having obtained electro-neutral S-trimers in boxes of water, a two-stage minimization was carried out with the steepest-descent algorithm for S-with-NAG; first with define=-dflexible option and then run without it (rigid water) and latter minimization for S-without-NAG. Following minimizations, each structure was equilibrated in two steps as 100 ps NVT and 100 ps NPT respectively with position constraints for the protein, to stabilize the temperature at 300 K and the pressure at 1 bar. Initial velocities were generated from a Maxwell–Boltzmann distribution at 300 K at the NVT. Production runs of MD simulations were carried out for 100 ns in the absence of position constraints. For this data collection part for the trajectory analysis, the temperature was kept constant with a coupling time constant of 0.1 ps using a velocity rescaling thermostat [40] with two coupling groups (Protein and non-Protein groups). The pressure was kept constant with a coupling time constant of 2.0 ps using a Berendsen barostat [41] with a coupling time constant of 2.0 ps. To examine the time evolution of the S trimers (S-without-NAG and S-with-NAG separately), Newton’s equation of motion was integrated numerically using the Leap-frog algorithm with a 2 fs time step. The linear constraint solver (LINCS) algorithm [42] was applied to bonds involving hydrogen atoms. Short-range electrostatic and van der Waals interactions were cut off at 1.4 nm using the Verlet scheme [43] and long-range electrostatic interactions were calculated using the particle mesh Ewald (PME) [44] algorithm. Long range dispersion corrections for both the energy and the pressure were applied to crank into the truncation of van der Waals terms. Periodic boundary conditions were applied in the *x, y* and *z* directions. All the calculations were carried out using the GROMACS 2019 [35] code and the GROMOS 54a7 force field [37] extended to include the ANX and NAN residues. The snapshots were obtained using the visual molecular dynamics (VMD) software [45].

### Results and Discussion

The coronavirus spike protein has a coating of N-linked glycans in various locations (see Table 1). To get the glycan bound to the S protein, we modified the Gromos 54a7 force field including ANX and NAN residues. Then, to understand if the NAG molecules lead the changes in the dynamics of the S protein, we analyzed the MD trajectories in the presence (S-with-NAG) and absence of NAG (S-without-NAG).

Structural stability along the simulations was analyzed using the backbone root-mean-square-deviation (RMSD) relative to the energy-minimized configuration of the starting structures in three different manners. First, the RMSD of the S-without-NAG and S-with-NAG were compared. The RMSD values for both systems range from 0.5 nm to 0.6 nm which are consistent with the results of [27] for the corresponding temperature; and it was observed that both systems reach stability after the first 20 ns of the simulation time (Figure 4a). Second, since the S protein is made up of two domains, S1 and S2, we also analyzed the RMSDs of these parts without and with NAG separately. The results are as follows.

- RMSD values of the S1 region is greater than that of the S2 region for both without and with NAG structures (Figure SI-1a and Figure SI-1b). Thus, it might be concluded that the S2 region is slightly more stable than the S1 region which is consistent with [**27**]
- For the S1 region (Figure 4b), after the 20 ns simulation, the *difference* in the RMSD values between S1-without-NAG and S1-with-NAG ranges from 0.04 nm to 0.09 nm and S1-with-NAG has greater RMSD values than the S1-without-NAG.
- In the S2 region (Figure 4c), the *difference* in the RMSD values between the S2-without-NAG and S2-with-NAG ranges from 0.02 nm to 0.07 nm and these differences are less than that of the S1 region. Moreover, unlike the S1 region (Figure 4c), S2-wihout-NAG has greater RMSD values than S2-with-NAG. It can be inferred that the presence of the NAG molecule makes the S2-domain more stable.

**Figure 4.**
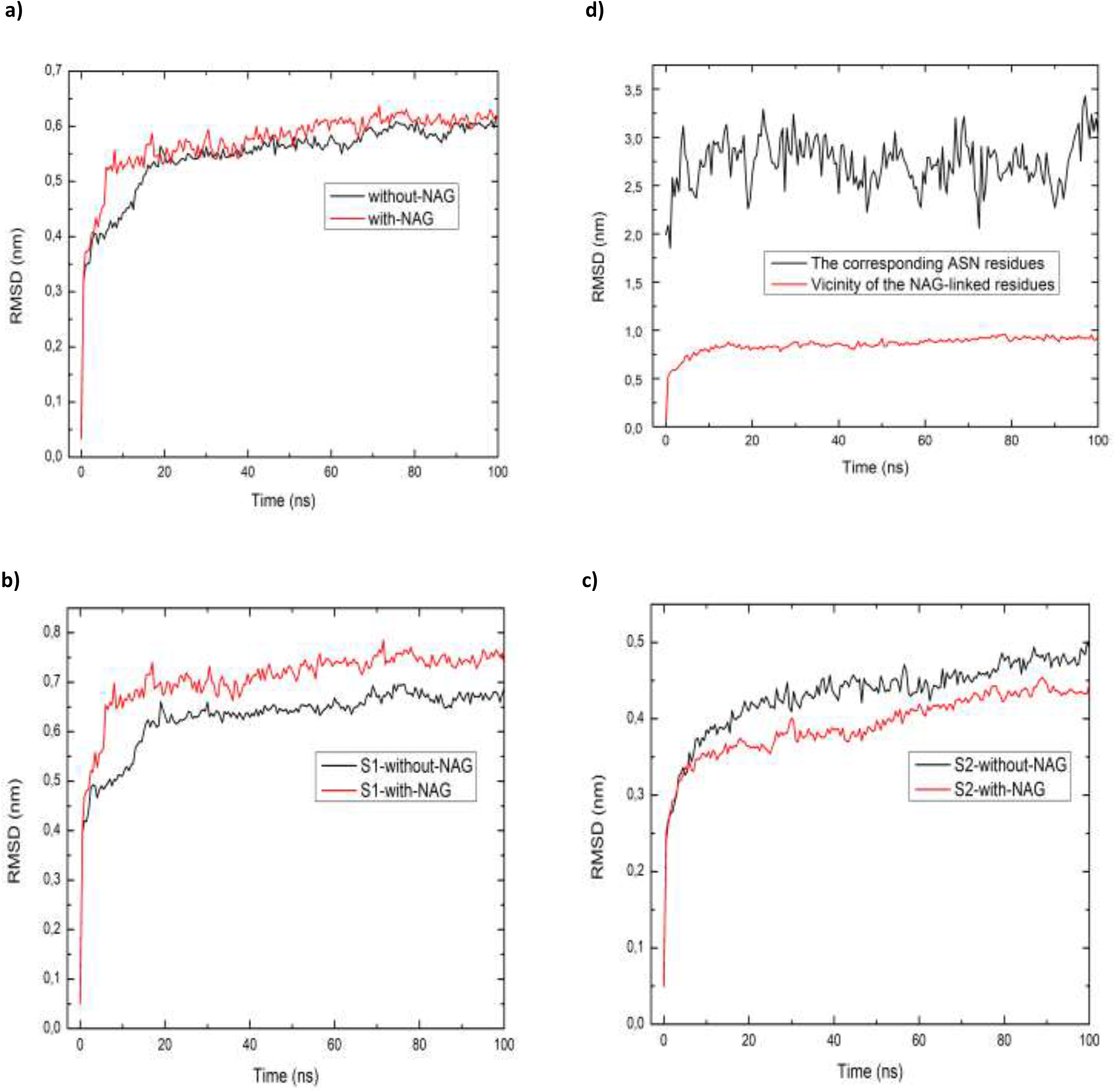
Comparison of the Root mean square deviation (RMSD) of the **a)** S-without-NAG and S-with-NAG **b)** S1 domain without and with NAG **c)** S2 domain without and with NAG **d)** Vicinity of NAG molecules

Third, since the NAG molecules are known to have important roles in the viral infection, we examined the vicinity of the NAG-linked residues and the corresponding ASN residues. Namely, we calculated the RMSD values between the NAG-linked amino acids (ANX and NAN; Table 1) and only amino acids (that is ASN aminoacids without NAG). Striking differences between them are observed; that is to say the RMSD value of NAG-linked amino acids, which is 0.9 nm and below, appears to be more stable than the one without NAG which is 2.7 nm on average (Figure 4d). It can be inferred that the N-linked glycans make the ASN amino acids more stable.

As a measure of compactness, that is to get an idea of the overall dimensions of the structures, the radius of gyration (Rg) was calculated throughout the simulations in three different ways. As in the case of RMSD results, gyration (Rg) calculation of S-without-NAG and S-with-NAG (Figure 5a) showed that the two structures exhibited nearly similar fluctuations. Gyration values of the S1 region is greater than the S2 region for both without and with NAG structures (Figure SI-2a and Figure SI-2b); 4.5 nm for S1 and 3.5 nm for S2 for both. S1 and S2 regions in S-with-NAG (Figure SI-2c) are larger than S1 and S2 regions in S-without-NAG (Figure SI-2d), as expected. However, if one zooms into the vicinity of the NAG-linked amino acids, one observes that the structure without NAG is more floppy; compact-uncompact, than the structure with NAG (Figure 5b).

**Figure 5:**
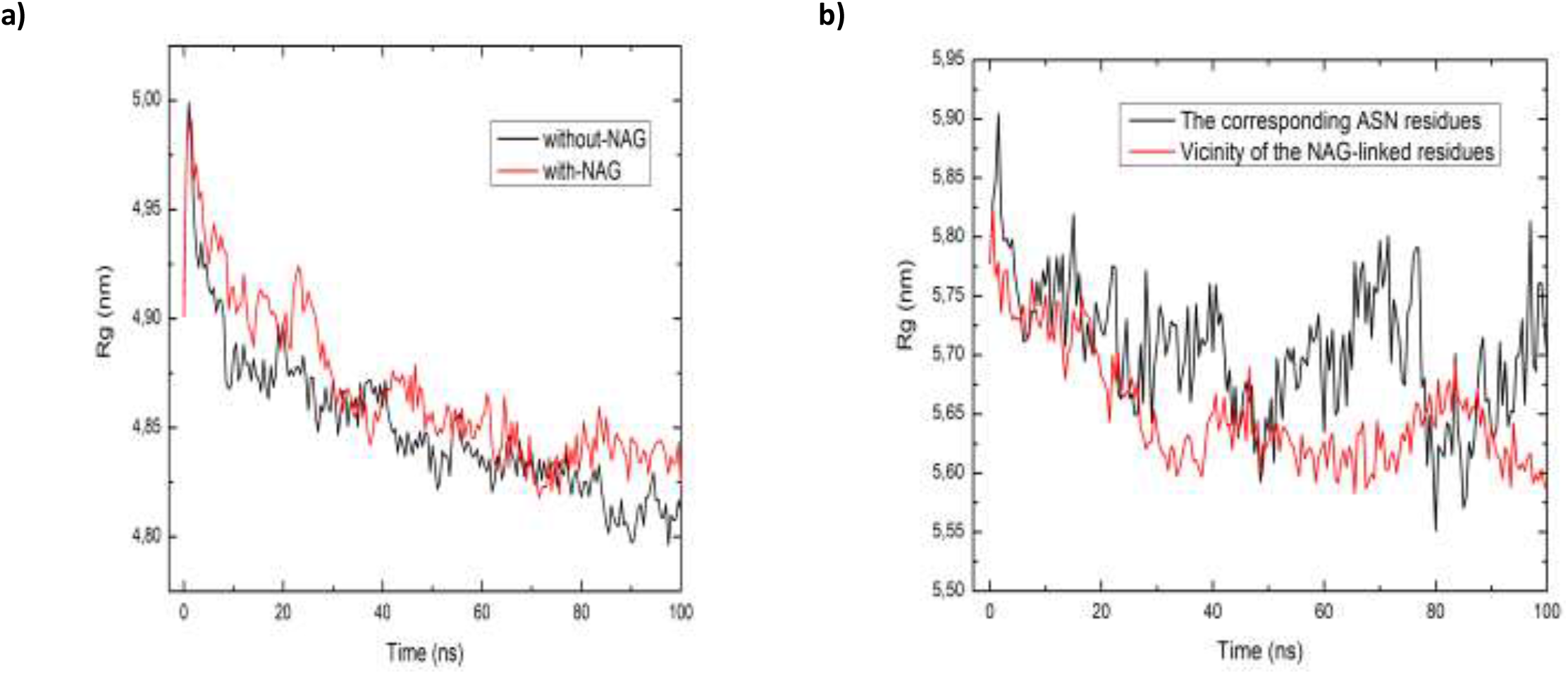
Comparison of Radius of gyration (Rg) plot of a) S-without-NAG and S-with-,NAG b) vicinity of the NAG molecules and the corresponding ASN residues.

Moreover, solvent accessible surface area (SASA) as a function of time was analyzed. SASA plot of S-without-NAG and S-with-NAG showed no considerable difference (Figure 6a); that is, there is no remarkable shielding by glycans (consistent with [9]). SASA of S1 is greater than S2 for both structures (Figures SI-3a and SI-3b). The presence of NAG did not cause much fluctuation in the SASA of the S1 region (Figure 6b), but reduced the fluctuation in S2 region (Figure 6c). SASA of NAG-linked amino acids (Figure 6d) are almost twice more than the SASA of the ASN aminoacids without NAG

**Figure 6:**
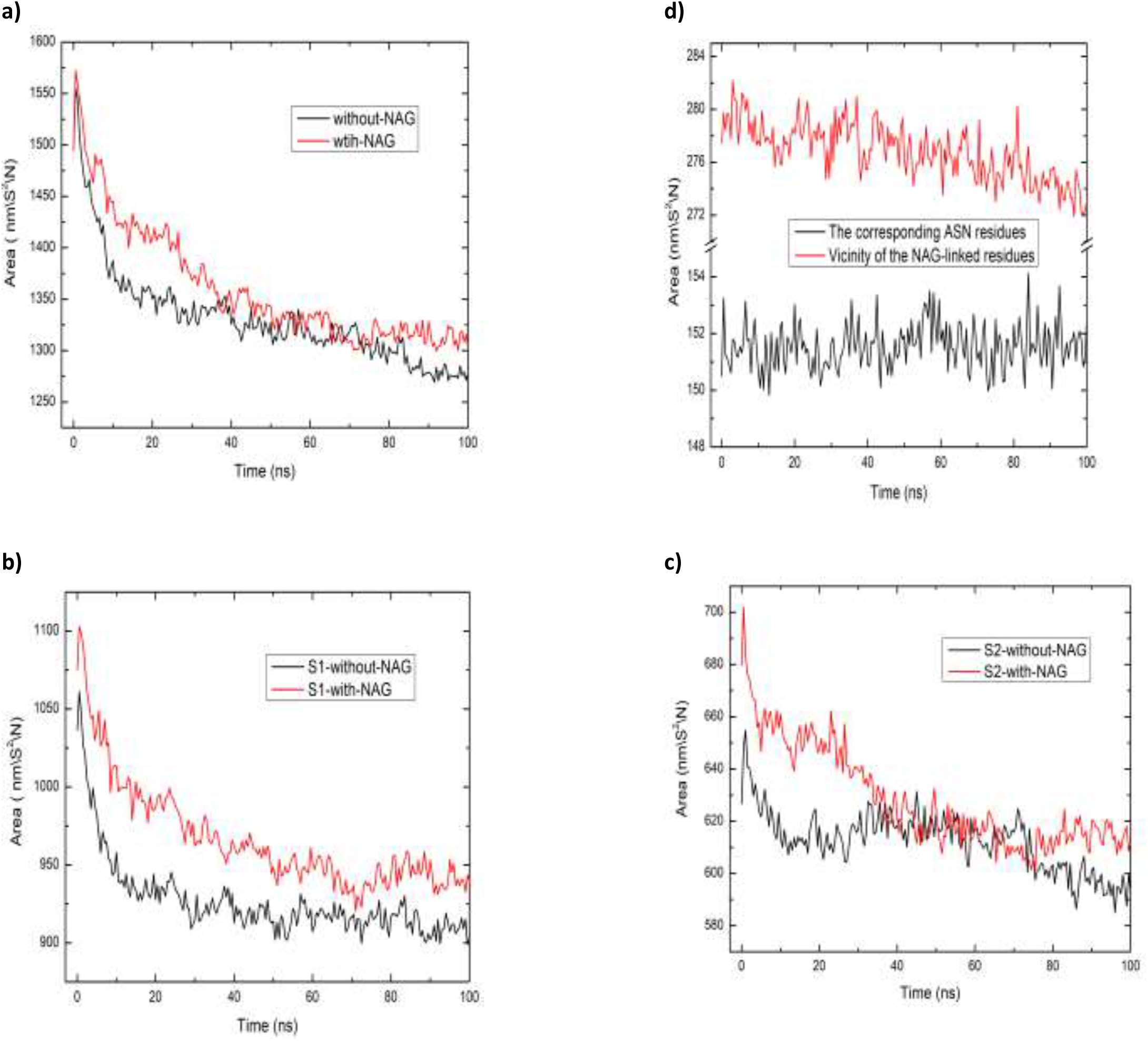
Comparison of solvent accessible surface area (SASA) of the a) S-without-NAG and S-with-NAG **b)** S1 domain without and with NAG **c)** S2 domain without and with NAG **d)** vicinity of the NAG molecules and the corresponding ASN residues.

We calculated the solvent accessibility of the individual glycan residues and the corresponding ASN residues. As seen from the Table 2 and Figure 6d, for all three-chains, residues ANX and NAN have almost 2 times more solvent-accessible surface areas than the corresponding ASN residues without NAG. Also, NAN has a larger solvent-accessible surface area compared to ANX.

**Table 2:** SASA of the individual glycan residues and corresponding ASN residues.

| Residue # | A-without-NAG | A-with-NAG | B-without-NAG | B-with-NAG | C-without-NAG | C-with-NAG |
| --- | --- | --- | --- | --- | --- | --- |
| 80 | 2.758 | 5.151 | 2.758 | 5.169 | 2.739 | 5.137 |
| 141 | 2.684 | 5.036 | 2.748 | 5.009 | 2.753 | 5.113 |
| 184 | 2.726 | 5.095 | 2.756 | 5.021 | 2.742 | 5.060 |
| 253* | 2.761 | 7.497 | 2.736 | 7.533 | 2.767 | 7.450 |
| 301 | 2.779 | 5.119 | 2.754 | 5.137 | 2.729 | 5.083 |
| 350 | 2.748 | 5.018 | 2.745 | 4.975 | 2.733 | 5.092 |
| 362 | 2.761 | 5.137 | 2.757 | 5.097 | 2.771 | 5.102 |
| 622 | 2.719 | 5.104 | 2.745 | 5.152 | 2.734 | 5.075 |
| 635 | 2.734 | 5.146 | 2.756 | 5.160 | 2.749 | 5.133 |
| 676 | 2.774 | 5.053 | 2.763 | 5.121 | 2.747 | 5.096 |
| 728 | 2.778 | 4.497 | 2.754 | 4.945 | 2.756 | 5.133 |
| 736 * | 2.731 | 7.397 | 2.744 | 7.470 | 2.768 | 6.675 |
| 820 * | 2.772 | 7.392 | 2.754 | 7.178 | 2.744 | 7.511 |
| 1093 | 2.782 | 5.093 | 2.748 | 4.903 | 2.776 | 5.139 |
| 1117 NAN | 2.707 | 7.371 | 2.687 | 7.127 | 2.732 | 6.632 |
| 1153 NAN | 2.739 | 7.402 | 2.744 | 7.423 | 2.763 | 7.442 |

In the case of hydrogen bonds (Figure 7) the cut-off distance and the cut-off angle criteria were set to 0.35 nm and 30° respectively. This analysis showed that the presence of the NAG leads to a decrease in the hydrogen bonds network established within the structure. This further supports the DSSP results (Figure 8a and 8b). Namely; in the S-with-NAG structure, the B-Sheet content decreases from 27% to 24%, while the coil content increases from 28% to 30%. The analysis of secondary structure as function of time was made using Dictionary of Secondary Structure of Proteins (DSSP) program by Kabsch and Sander [46]. There are no significant differences among the number of hydrogen bonds between the regions without and with NAG structures (Figure SI-4a, Figure SI-4b, Figure SI-4c)

**Figure 7:**
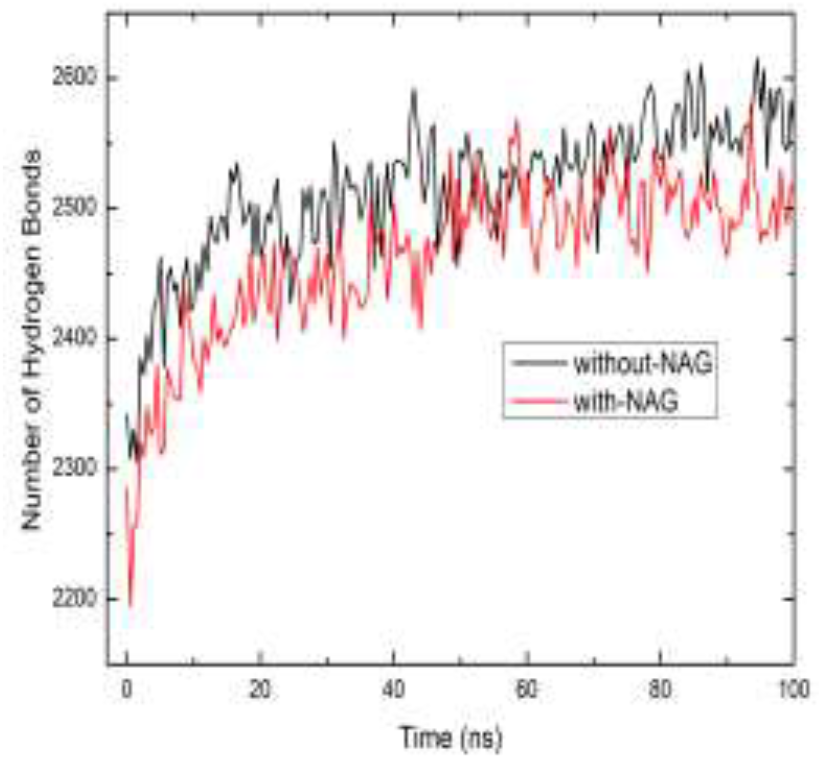
Intra protein hydrogen bond pattern of S-without-NAG and S-with-NAG

**Figure 8:**
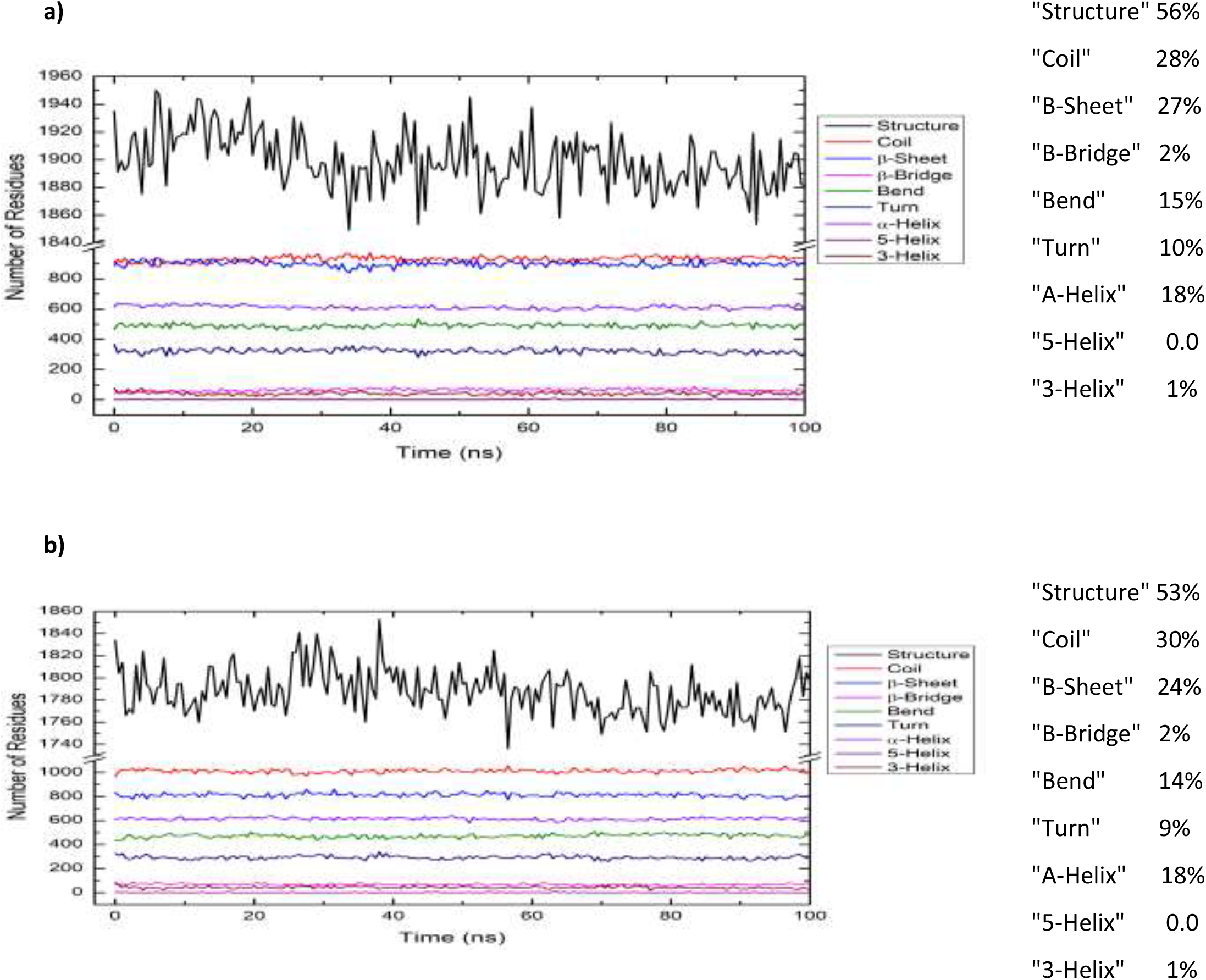
Secondary Structure content of the a) S-without-NAG b) S-with-NAG

## Conclusions

The spike glycoprotein is the trimer structure decorated with the NAG molecules, located at the outermost part of the SARS-CoV-2, which enters the respiratory system by interacting with the human receptor ACE2. Some experiments and theoretical simulations show the importance of the NAG molecule on the stability of the different molecules. At the molecular level, however, the effect of NAG molecules on the SARS-CoV 2 structure is still not clear. In this study, 100 ns MD simulations of closed SARS-CoV-2 S (6VXX.pdb) were performed without and with NAG to better understand the structures in atomic details and the importance of NAG molecules on the stability of the SARS-CoV-2 S glycoprotein. This comparative study revealed that even though there are no striking differences between the stability of the full structures in the absence or presence of the NAG molecules, glycans increase the local stability; that is to say the vicinity of NAG molecules are much more stable than the corresponding ASN residues. Moreover, the S1 domain is more flexible than the S2 domain. Overall, these results can enlighten the nature of the SARS-CoV-2 providing an information at the atomic level, as well as be useful for vaccines and therapeutics development.

## Acknowledge

The numerical calculations reported in this paper were fully/partially performed at TUBITAK ULAKBIM, High Performance and Grid Computing Center (TRUBA resources).

