## Supplementary Information for "Investigation of the Effects of N-Linked Glycans on the Stability of the Spike Protein in SARS-CoV-2 by Molecular Dynamics Simulations"

a) b)

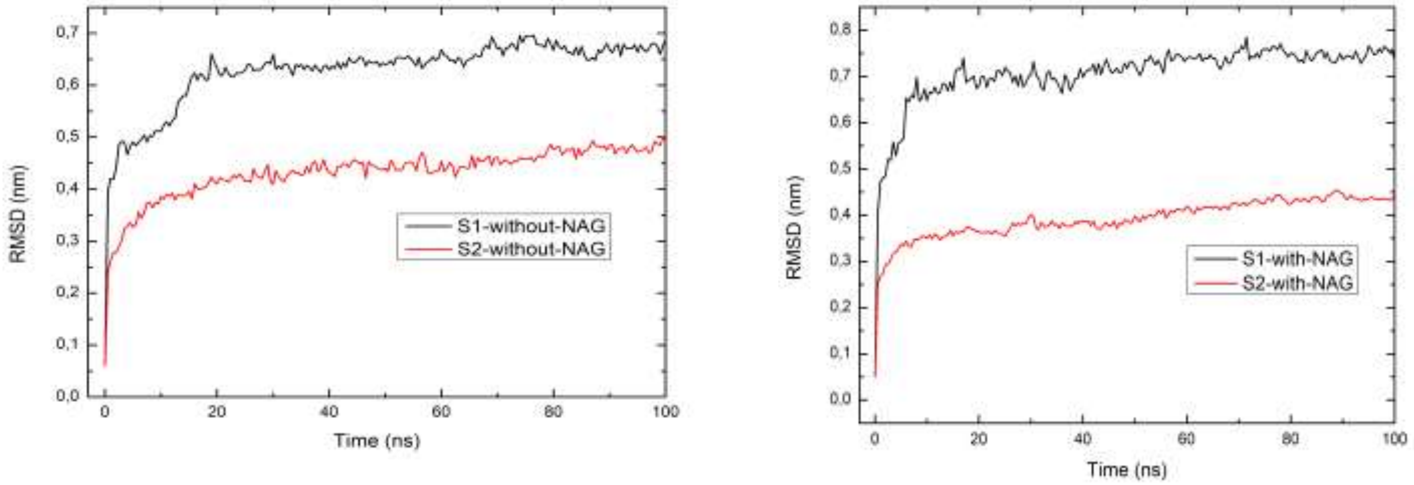


**Figure SI-1:** Comparison of the root mean square deviation (RMSD) of the **a)** S1 and S2 domains without NAG **b)** S1 and S2 domains with NAG

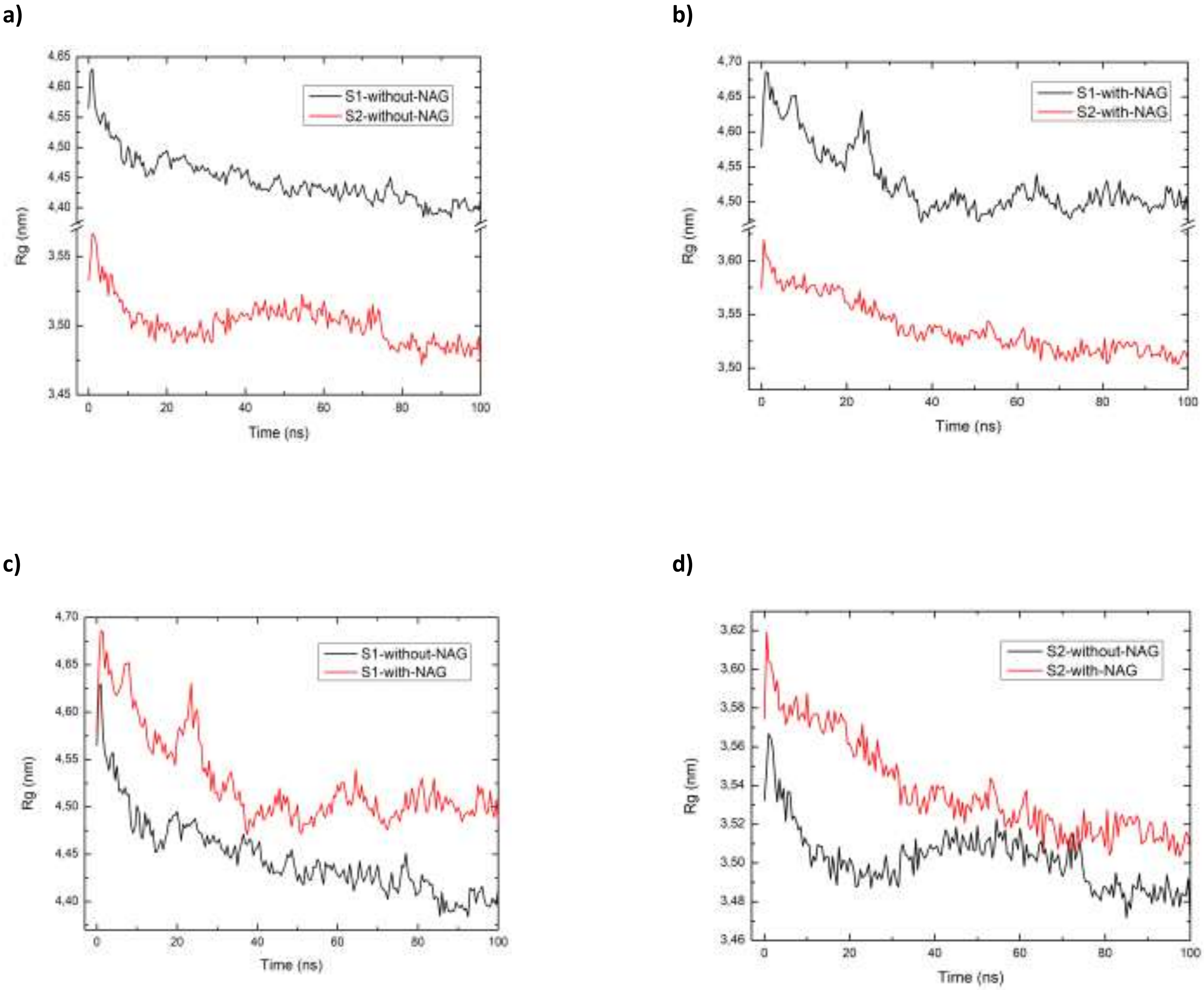


**Figure SI-2**: Comparison of Radius of gyration (Rg) plot of a) S1 and S2 somains with NAG b) S1 and S2 domains with NAG c) S1 domain without and with NAG d) S2 domain without and with NAG.

a) b)

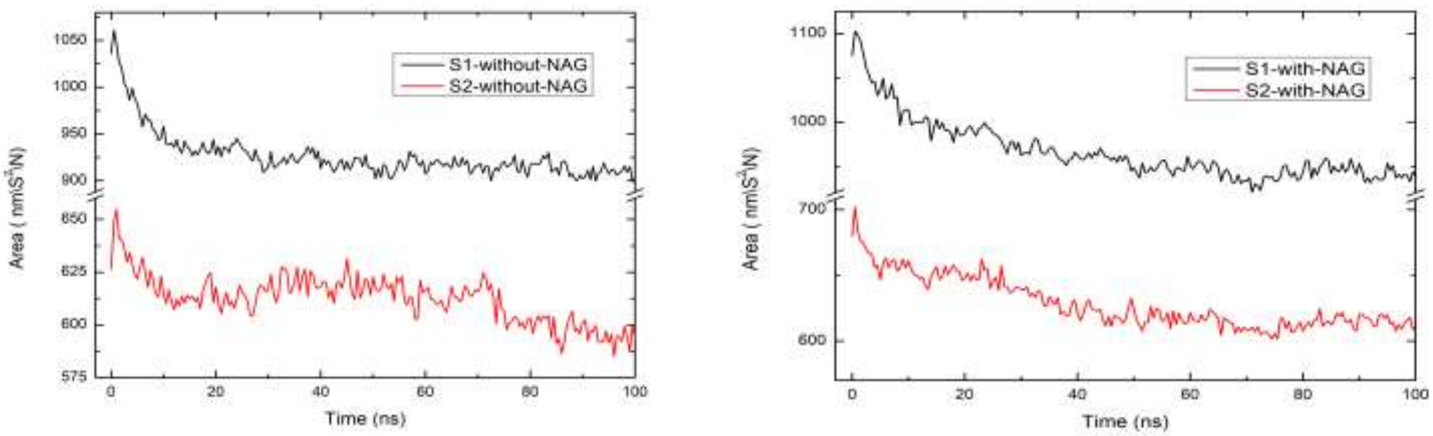


**Figure SI-3:** Comparison of solvent accessible surface area (SASA) of the a) S1 and S2 domains without NAG b) S1 and S2 domains with NAG.

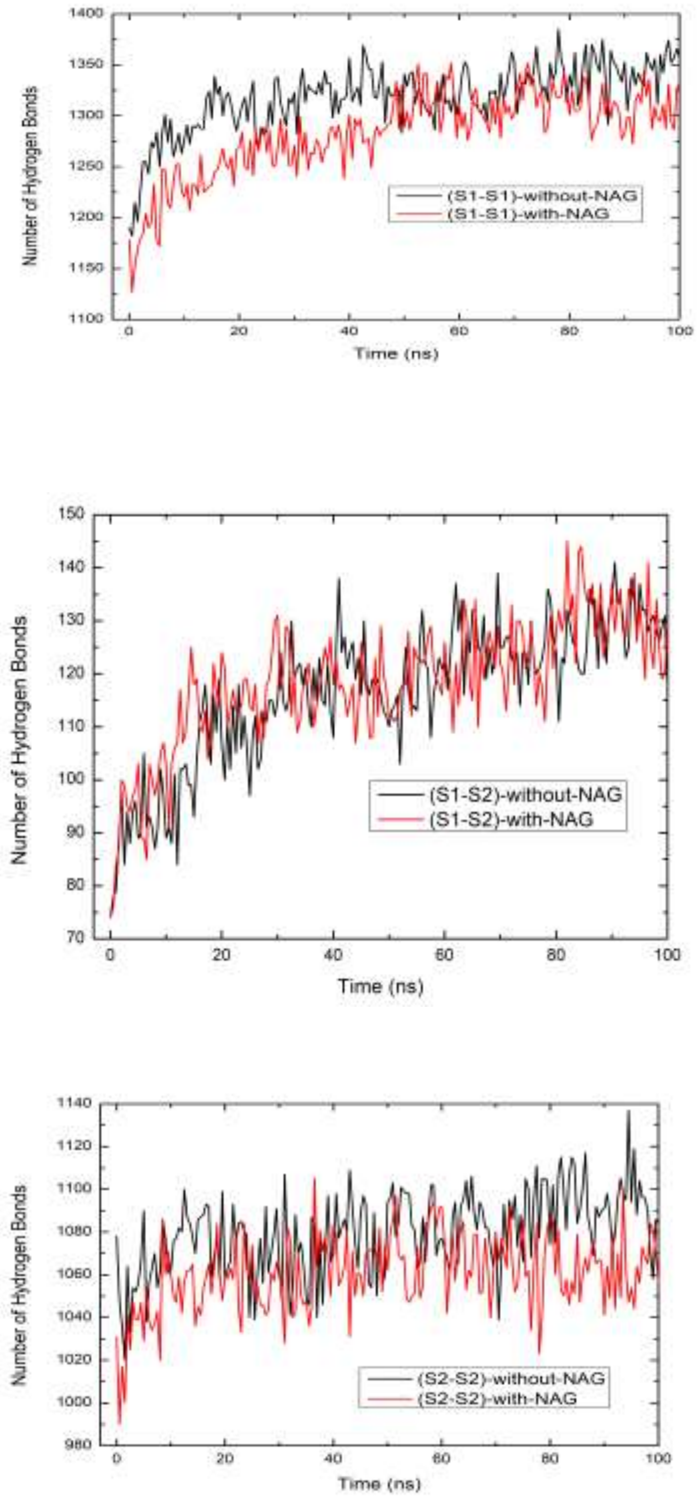


**Figure SI-4:** Comparison of number of intra protein hydrogen bond between a) S1-S1 with and without NAG b) S1-S2 with and without NAG c) S2-S2 with and without NAG.
